# Continuous theta-burst stimulation of the ventromedial prefrontal cortex reduces Pavlovian go-invigoration and enhances thalamo-striatal reward prediction error signals

**DOI:** 10.1101/2025.11.11.686520

**Authors:** Hans-Christoph Aster, Jasmin Schmitt, Martin J. Herrmann, Daniel Zeller, György Homola, Thomas Kampf, Mirko Pham, Sebastian Walther, Marcel Romanos, Vanessa Scholz, Lorenz Deserno

## Abstract

The ventromedial prefrontal cortex (vmPFC) represents value, i.e. whether environmental cues are good or bad. Rewarding cue value is aligned with a tendency to act or to inhibit in case of potential punishment, a so-called Pavlovian Bias. Value-related vmPFC activation may therefore impact Pavlovian bias. However, it remains unknown how the activity in the vmPFC modulates the Pavlovian bias. Thus, we applied continuous theta-burst stimulation (cTBS), a non-invasive protocol that transiently reduces cortical excitability, over the vmPFC in a single-blinded, between-subject study. 90 healthy adults performed a motivational Go/NoGo learning task during fMRI after receiving cTBS (n=45) or sham (n=45) stimulation. Behaviourally, cTBS reduced overall Go response rates and slowed reaction times. Computational modelling showed a decrease in the learning rate, selectively for rewarding outcomes. Computational fMRI analysis showed stronger reward prediction error (RPE) signals in the vmPFC, mediodorsal thalamus, and left dorsal striatum after cTBS, without changes in neural signals at cue onset. These findings provide causal evidence that the vmPFC drives the action-invigoration component of the Pavlovian bias.

**Significance statement:** We causally link ventromedial prefrontal cortex (vmPFC) function to motivated action in humans. In a single-blind cTBS-fMRI study (n=90), transient vmPFC inhibition reduced Go responding, slowed reaction times, and selectively lowered the positive learning rate early in learning. At the same time, reward-prediction-error signals increased in mediodorsal thalamus, dorsal striatum, and vmPFC, while cue-onset value/choice signals were unchanged. Together, the results indicate that vmPFC normally speeds value updating and invigorates actions; when inhibited, control shifts toward slower, subcortically supported learning under higher uncertainty.

## Introduction

Due to limited resources, the brain often relies on biases, such as the Pavlovian bias, which is the tendency to act when rewards are expected and to withhold action when punishment is anticipated (Guitart-Masip et al., 2014a; Huys et al., 2015). This bias can be overridden by a flexible instrumental system that acquires accurate action-outcome contingencies (Rangel et al., 2008). Conflicts emerge when instrumental learning requires responses incongruent with this bias (Rescorla and Solomon, 1967; Gershman et al., 2021). The Pavlovian bias is typically assessed with the motivational Go/NoGo task, where action and cue valence are orthogonalized (Guitart-Masip et al., 2012b; van Wouwe et al., 2015).

Pavlovian bias is thought of as an evolutionarily conserved (“hard-wired”) mechanism of rather fast and automatic behavioural control (Raab and Hartley, 2020). According to Frank and colleagues’ neurobiological computational model, transient dopamine increases (“bursts”) from the midbrain to striatum, following positive reward predictions, transiently boost activity of D1-neurons in the direct pathway, disinhibiting thalamo-cortical drive and biasing the organism toward “Go” responses. Meanwhile, transient decreases (“dips”) following negative predictions activate D2-expressing neurons in the indirect pathway, thereby heightening pallido-thalamic inhibition and promoting “NoGo” responses (Maia and Frank, 2011; Collins and Frank, 2014). Yet, dopamine signalling within the striatum alone is unlikely to fully account for the neural basis of Pavlovian biases. For example, and in contrast with predictions based on striatal learning, acute pharmacological enhancement of dopamine transmission with levodopa reduced the computational Pavlovian parameter and increased instrumental learning (Guitart-Masip et al., 2014b). A recent PET study found no direct association between striatal dopamine synthesis capacity and Pavlovian bias. fMRI data with this task indicates that striatal activity primarily encodes instrumental actions (Go vs. NoGo) rather than cue valence (Win vs. Avoid) (Guitart-Masip et al., 2011; Guitart-Masip et al., 2012b). Therefore, cue-valence encoding may be a key contributor to the Pavlovian bias. The ventromedial prefrontal cortex (vmPFC), among other cortical regions, encodes cue valence (Gershman et al., 2021; Algermissen et al., 2024). Enhanced vmPFC cue-valence encoding may modulate phasic striatal dopamine signals through the vmPFC’s extensive connections with the ventral striatum, thalamus, and amygdala (Haber and Calzavara, 2009; Hiser and Koenigs, 2018). Interestingly, recent work using combined fMRI-EEG emphasizes a role for the representation of cue valence signals in cortical regions, including vmPFC, that temporally preceded and shaped biased PE in the striatum (Cavanagh et al., 2013; Swart et al., 2018; Algermissen et al., 2021).

Despite this accumulating correlational evidence, work that involves a direct manipulation of vmPFC activity while assessing its influence on Pavlovian bias is still missing (Algermissen et al., 2021; Messimeris et al., 2023). Thus, to manipulate the contribution of the vmPFC in this regard, we used continuous theta-burst stimulation (cTBS) over the vmPFC, which transiently reduces cortical excitability for 30–60 minutes (Huang et al., 2005; Wischnewski and Schutter, 2015). The main goal of this study was to experimentally test how inducing a temporary ‘virtual lesion’ in the vmPFC using cTBS would affect the behavioural expression of Pavlovian bias and its neural correlates during the motivational Go/NoGo learning task. At the neural level, we expected transient vmPFC inhibition via cTBS to reduce cue-valence–related BOLD responses in this region. Further, this may result in enhanced outcome-related activity in the striatum, potentially reflecting increased reliance on striatum-mediated learning when vmPFC-derived valence signals are reduced. At the neural level, we expected transient vmPFC inhibition via cTBS to reduce the BOLD signal typically observed in this region when win as compared to avoid cues presented. Behaviourally, we hypothesized that cTBS-induced blunting of cue valence signals would attenuate the behavioural strength of Pavlovian bias. Results from this study may not only elucidate the role of the vmPFC in Pavlovian Bias but could also provide critical insights for targeted therapeutic interventions using cTBS for disorders associated with altered Pavlovian bias, such as addiction (Chen et al., 2023a) and obsessive-compulsive disorder (Peng et al., 2022).

## Methods

### Study design

For this study, 90 subjects aged between 18 and 40 were recruited through public advertising. Inclusion criteria for the study were a negative SCID screening interview (Wittchen et al., 1997), which tests for the presence of mental disorders in the present or past, and sufficient knowledge of German to understand the task and study instructions. Exclusion criteria were contraindications to TMS or MRI (such as metal on or in the body, implants, or claustrophobia), the presence of internal, neurological, psychiatric, or other serious medical illness, and the presumption of pregnancy. At the start of the study, we randomly assigned subjects to a cTBS or a Sham-cTBS group. Throughout the recruitment process, subjects were assigned to the groups such that the groups were matched in age and gender.

In a single-blinded, between-subjects design, subjects were invited to a study appointment, in which they received either an active cTBS or a Sham TMS. A between-subjects design was chosen due to the significant irritation of the sensory skin nerves induced by a cTBS (Zhao et al., 2020). Consequently, a within-subject design with two study appointments would have automatically caused unblinding of the subjects regarding group assignment. Further, a within-subjects design may be prone to strong repetition or training effects (Scholz et al., 2022; Scholz et al., 2025). At the beginning of the study session, subjects were instructed about study procedures. Thereafter, depending on group assignment, they either received cTBS or Sham-cTBS for approximately 40 seconds and were then immediately transferred to the MRI scanner to complete the motivational Go/NoGo task (citation). In total, the subjects completed two rounds of either cTBS or Sham-cTBS, immediately followed by two behavioural tasks in the fMRI, see figure 1c. The second behavioural task is not part of this publication and will be reported elsewhere. The order of the two behavioural tasks was counterbalanced. Subjects were continuously surveyed throughout the study regarding their mood, tension, and fatigue on a scale of −5 to 5 (before the initial cTBS/Sham session, after the first cTBS/Sham session, after the initial MRI experiment, after the second cTBS/Sham session, and finally after the second MRI experiment). The Ethics Committee of the Medical Faculty of Würzburg gave its positive vote for this study (file number 9-21).

**Figure 1.**
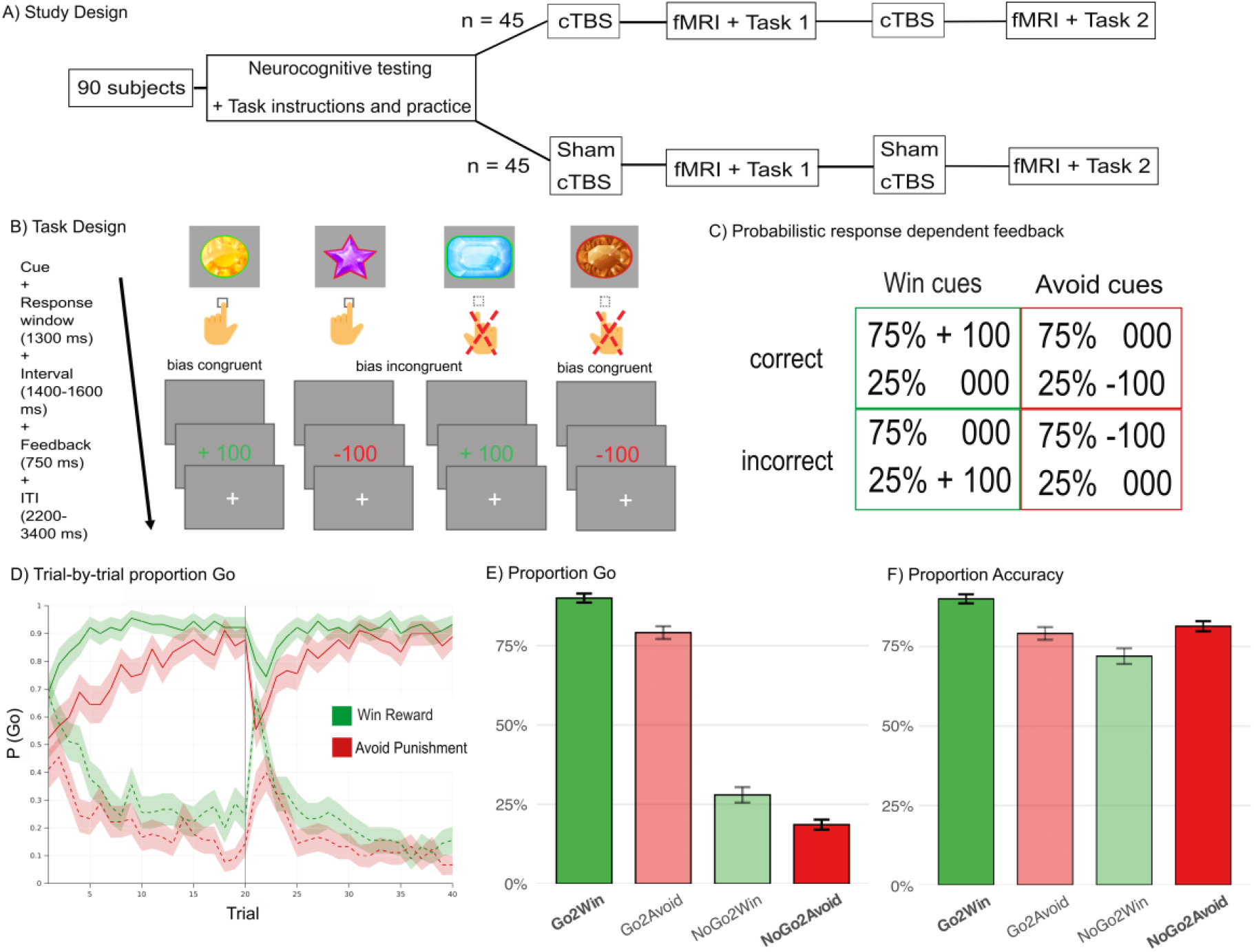
A) Timeline of the study procedure. As the cTBS effect was expected to last around 30 minutes, cTBS/Sham-CTBS was repeated after participants finished the first task. B) Trial structure and cue types. The motivational Go/NoGo task consisted of four trial types based on cue-action-outcome associations: Go-to-Win, Go-to-Avoid, NoGo-to-Win, and NoGo-to-Avoid. In bias-congruent conditions (Go-to-Win, NoGo-to-Avoid), the required action aligns with the Pavlovian bias (i.e., approach in anticipation of reward, inhibition in the context of punishment). In contrast, bias-incongruent conditions (Go-to-Avoid, NoGo-to-Win) demand responses that conflict with these Pavlovian tendencies and are typically more difficult to execute. Each trial began with the presentation of a cue (1300 ms), during which participants chose to either respond via button press (Go) or to withhold a response (NoGo). Subsequently, an outcome (reward, punishment, or neutral feedback) was displayed for 750 ms. The inter-trial interval (ITI) varied randomly between 2200 and 3400 ms in 200 ms increments. C) Feedback contingencies. Each cue was probabilistically associated with a correct outcome, using a 75%/25% feedback contingency to enable learning while maintaining some uncertainty. D) Trial-wise choice behavior. The plot shows the evolving probability of executing a Go response (P(Go); ± Standard Error), computed using a sliding window of 5 trials. Solid lines represent Go-cue trials, dashed lines indicate NoGo-cue trials, and data are collapsed across both groups. A pronounced Pavlovian bias is evident from the beginning of the experiment, with higher Go response probabilities for reward-predicting cues compared to punishment-predicting cues that require inhibition. Throughout the experiment, participants increasingly select the correct Go response for Go cues over time, reflecting learning. E) Cue-specific Go response probability. Average P(Go) across trials is shown separately for each cue type. Bias congruent cue types are printed in bold. Pavlovian biases are evident in the elevated P(Go) for reward cues, even when Go is not the optimal action. F) Average accuracy across trials is shown separately for each cue type.

### TMS protocol

The cTBS group was stimulated using a MagPro X100 stimulator (Medtronic A/S, Skovlunde, Denmark) and a MagVenture MC-B70 figure-of-eight coil (MagVenture A/S, Farum, Denmark). In the Sham-TMS group, the same setup was used but with an MC-P-B70 placebo coil. This placebo coil has a magnetic field shielding that produces a field strength reduction of 80%. The number of stimulations during the Sham stimulation corresponded to that of the MC-B70 coil during cTBS. As a stimulation point for the vmPFC, we chose the electrode position FPz according to the international 10-20 system, as used in previous studies (Herrmann et al., 2017; Manuel et al., 2019; Dantas et al., 2023). For the cTBS, 600 stimuli in total were applied at 80% active motor threshold (AMT) during an uninterrupted train of 40 seconds (Huang et al., 2005; Wittmann et al., 2021). The coil handle was directed caudally as the coil was placed tangentially to the forehead above the FpZ localization. The AMT was determined as the minimum intensity of stimulation required over the left M1 to elicit motor-evoked potentials (MEPs) with amplitudes of at least 50 µV in at least 5 out of 10 trial stimulations during voluntary muscle activation of the right first interosseous muscle at about 20% of maximal force. Electromyography (EMG) data were recorded from the right first interosseous muscle using surface electrodes (Odorfer et al., 2019).

### Sample characterization

To detect possible group differences in personality traits or neurocognitive parameters potentially influencing task performance, participants completed the following online questionnaires in advance and completed neurocognitive testing before the start of the experiments. First, participants also reported their vigilance (ranging from −5 to 5) before and after the experiment, as vigilance has been shown to influence performance in Go/NoGo tasks (Pershin et al., 2023), as well as their assumed group assignment afterward to assess blinding integrity and potential expectancy effects. Clinical questionnaires included the Beck Depression Inventory II (Jackson-Koku, 2016), the State-Trait Anxiety Inventory (Tenenbaum et al., 1985), the Barratt Impulsiveness Scale (BIS-15, (Stanford et al., 2009)), the Obsessive-Compulsive Inventory (Revised OCI-R, (Gönner et al., 2007)) and an ADHD symptom questionnaire (ADHD Selbstbeurteilungsskala, (Retz et al., 2013)). To test general cognitive skills such as processing speed or working memory, subjects completed the trail-making task (Tombaugh, 2004), the listening span task (Daneman and Carpenter, 1980; Swart et al., 2017), and the digit span task (Blackburn and Benton, 1957).

### Motivational Go/NoGo task

We employed a well-established motivational Go NoGo task capturing the impact of motivational biases on choice behaviour (van Nuland et al., 2020; Scholz et al., 2022). In this task, participants are shown different cues (gems in different colours and shapes) for 1300 milliseconds (ms), after which they must decide whether to respond with a Go response (pressing the response handle in the right hand) or not (NoGo response). Immediately after the disappearance of the cue, subjects received feedback (cue-feedback interval between 1400 and 2600 ms, steps of 200ms) for 750 ms in the centre of the screen. Valence was instructed during practice trials, such that cues with a green edge (win) were followed by reward (100 points) or neutral feedback (0 points), whereas cues with a red edge (avoid punishment cues) were followed by neutral feedback (0 points) or punishment (−100 points). Thus, depending on the trial, participants played to win points (green edge: Go2Win or NoGo2Win) or to avoid losing points (red edge: Go2Avoid or NoGo2Avoid). Throughout the course of the experiment, subjects had to learn which action resulted in the most wins or the least losses for the different cues, guided by the provided feedback. Since feedback was probabilistic (75/25), when a correct answer was given, a win appeared in 75% of the win-trials (+100 points), and neutral feedback was given in 75% of the avoid-trials (0 points). The purpose of the experiment was to collect as many points as possible at the end of the game. The task had two blocks, each with 80 trials. Each block had a new cue set and lasted 9 min (Total: 160 trials; 20 trials per cue per block, inter-trial intervals between 2200-3400 ms in steps of 200ms). To verify that participants understood the task, they played a practice round of 32 practice trials on a computer outside the MRI before completing the task in the scanner, and had to explain it to the instructor after the practice trials.

### Behavioural analysis

To assess the effects of the TMS intervention on Pavlovian biases on choice behaviour, we examined whether the probability of making a Go response, P(Go), changed according to the within-subject factors like cue valence (Win vs. Avoid Punishment, coded 1 and −1) and required action (Go vs. NoGo response, coded 1 and −1). Moreover, the model included intervention group (cTBS vs. SHAM, coded 1 and −1) as a between-group factor. To control for group differences in vigilance and the accuracy of the participants’ assumed group assignment, we included mean vigilance (standardized) and the accuracy of the assumed group assignment (coded 1 and −1) as fixed effects in our models. Our main interest was on effects involving the cTBS intervention with a specific focus on the 2-way interaction valence x TMS to determine whether group association affected the strength of Pavlovian biases in choice behaviour. We used the glmer function from the lme4 package in R (optimizer: bobyqa) to fit the following trial-by-trial generalized linear mixed models (GLMMs):

1. P(Go) ∼ Group* Action * Valence * Block * (Accuracy of group assignment + Vigilance) + (1 + Action * Valence * Block |subject),
2. Accuracy ∼ Group * Action * Valence * Block * (Accuracy of group assignment + Vigilance) + (1 + Action * Valence * Block |subject),
3. Reaction Time ∼ Group * Action * Valence * Block * (Accuracy of group assignment + Vigilance) + (1|subject).

For both the Go-Response and Accuracy models, we used a binomial distribution with a logit link function, including random intercepts and random slopes (with correlations). In contrast, for the Reaction Time (RT) model, we used a gamma distribution with an identity link function. This selection was informed by previous findings, such as those by Lo and Andrews (Lo and Andrews, 2015), demonstrating that this configuration aligns most effectively with RT data characteristics. Because the full random-effects structure for the RT model did not converge, we reduced it to only include random intercepts (i.e., no correlated slopes). This follows the general recommendation by Barr et al. (Barr et al., 2013). The maximal number of iterations was set to n = 1e+9. Results were considered statistically significant with p-values < 0.05.

### Computational modelling

To investigate possible computational mechanisms underlying the behavioural effects, we fitted a series of nested hierarchical reinforcement learning (RL) models (Swart et al., 2017; Scholz et al., 2022; Scholz et al., 2024). The model space included 7 models overall. For model fitting and subsequent model comparison, we used the CBM toolbox implemented in MATLAB (Piray et al., 2019), which relies on Hierarchical Bayesian Inference (HBI) and treats not only parameters but also models as random effects. To account for potential block effects, model parameters were estimated separately for each task block.

The initial model (M1) was based on a basic Rescorla-Wagner model (Rescorla and Wagner, 1972). It comprised a parameter for feedback sensitivity (ρ) and a learning rate (ε), allowing the model to learn and update the value of each action (a: Go / NoGo) for each stimulus (s) on each trial t.

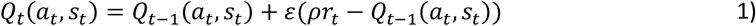

The subsequent model (M2) introduced a third parameter known as the ‘Go bias’ (b), embodying the general tendency to favour ‘Go’ responses. Model M3 then expanded on M2 by adding a motivational bias parameter (π). This parameter effectively captured the inclination to produce more ‘Go’ responses for cues associated with winning outcomes compared to cues associated with avoiding negative outcomes. These inherent biases were then integrated with the acquired Q values into the action weights (w):

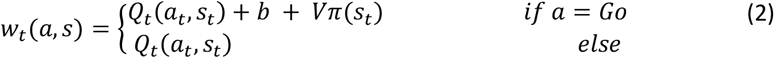

Here, V denoted the valence of cues (V_win_ = + 0.5; V_avoid_ = −0.5). Consequently, a positive value of π would amplify the action weight linked to ‘Go’ responses to cues associated with winning, while simultaneously reducing the weight for ‘Go’ responses to cues associated with avoiding negative outcomes. Concluding this process, the action weights underwent a transformation, converting them into actionable probabilities using a well-defined softmax function:

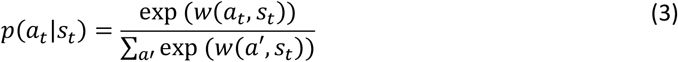

To probe whether the impact of feedback varies with motivational context or outcome valence, we extended the model space in four consecutive, nested steps. In Model M4, feedback sensitivity was allowed to differ between win and avoid cues (ρ_win_, ρ_avoid_) while keeping a single learning rate ε. Model M5 comprised one feedback-sensitivity parameter ρ but introduced separate learning rates for win (ε_win_) and avoid (ε_avoid_) trials. Model M6 partitioned feedback sensitivity by valence, estimating ρ_+FB for positive and ρ_-FB for negative feedback and a single ε. Finally, Model M7 retained a global ρ and estimated valence-specific learning rates (ε_+FB_, ε_-FB_), allowing gains and losses to drive learning at different rates.

To determine the model best suited to describe our empirical data, we used random effects model comparison (Stephan et al., 2009), computing the Laplace approximation of model evidence based on the individual level (Kass and Raftery, 1995; Daw et al., 2011). Subsequently, the group model evidence is derived to establish the “winning” model. Model evidence was assessed using model frequency and protected exceedance probability (PXP) (Piray et al., 2019), with the latter evaluating the most frequently expressed model (Stephan et al., 2009) while accounting for the possibility of chance results. To conclude the analysis, we then extracted hierarchical model parameter estimates from the winning model and compared group differences across model parameters between the TMS vs. Sham group using generalized linear mixed models (Parameter ∼ Group * Block + (1 + Block |subject)). Results were considered significant with a p-value < .05.

### MRI: data collection and preprocessing

Imaging data were collected using a 3 Tesla Siemens Prisma scanner and a 64-channel head coil. T1w images were acquired using a magnetization prepared rapid gradient echo (MPRAGE) sequence (TE = 3.17ms, TR = 2.4s, Inversion Time = 1s, 1×1×1mm voxel size). Functional data were collected using an echo-planar imaging sequence (TE = 22ms, TR = 2s, Flip angle = 80°, and 2.5×2.5×2.5mm voxel size). Field maps were also acquired to correct for inhomogeneity in the magnetic field (TE 1 = 5.19ms, TE 2 = 7.65ms, TR = 695ms, flip angle = 60°).

All data were pre-processed using SPM12 (Penny et al., 2011) running under MATLAB R2022b. Volumes were first corrected for slice-acquisition timing. Dual-echo gradient-echo field maps were transformed into voxel-displacement maps with the SPM FieldMap toolbox and applied during a single Realign & Unwarp step that simultaneously estimated rigid-body motion and corrected for susceptibility-by-movement interactions using 4th-order B-spline interpolation. The mean unwarped EPI was then coregistered to the individual T1-weighted MPRAGE. Anatomical images were segmented with the unified segmentation algorithm (Ashburner and Friston, 2005), and the resulting forward deformation fields were applied to all motion-corrected EPIs, resampling them to 2 × 2 × 2 mm and normalizing them into MNI 152 space. Finally, normalised images were smoothed with an isotropic Gaussian kernel of 6 mm full width at half maximum. The six head-motion parameters were carried forward as nuisance regressors in the first-level GLM.

### MRI: statistical analysis

First level: For each participant, a subject-specific general linear model (GLM) was estimated using SPM12 on the pre-processed and smoothed functional images. Five regressors of interest were included: Cue onset (collapsed across all four cue types), trial-wise model-derived choice certainty, entered as a parametric modulator at cue onset, button-press onset, outcome onset, and trial-wise model-derived reward-prediction error (RPE), entered as a parametric modulator at outcome onset.

All regressors were convolved with the canonical hemodynamic response function (HRF) implemented in SPM, without inclusion of temporal or dispersion derivatives. Six realignment parameters (three translations, three rotations) obtained during pre-processing were included as nuisance regressors. Single-subject contrast images were computed separately for each regressor of interest (cue onset, choice certainty, button press, outcome onset, and prediction error). To differentiate win versus avoid cues, we also fitted a model with separate cue onsets.

Second-level analyses were conducted separately for each of the first-level contrasts. Individual contrast images from each participant were entered into group-level analyses in SPM12. After excluding three participants due to pre-processing errors (failed normalization), the final sample for the fMRI analysis consisted of 87 individuals (44 cTBS, 43 Sham). Between-group comparisons (TMS > Sham and Sham > TMS) were tested using independent two-sample t-tests, while the main effects across all participants were assessed using one-sample t-tests. Whole-brain statistical inference was performed using a voxel-level threshold of p < 0.001 (uncorrected) combined with cluster-level family-wise error correction (FWE) at p < 0.05. Beyond the whole-brain contrasts, we carried out hypothesis-driven ROI analyses focused on the ventromedial prefrontal cortex (vmPFC) and the bilateral striatum. The vmPFC was selected because it served as the direct site of non-invasive stimulation in the present experiment, while both the vmPFC and striatum are well established as key nodes for the representation of subjective value (Bartra et al., 2013). ROIs were defined anatomically using labels from the Automated Anatomical Labeling (AAL) atlas (Tzourio-Mazoyer et al., 2002). The striatal ROI was generated by combining the atlas masks for the caudate and putamen, which together encompass the nucleus accumbens (not separately delineated in the AAL). The vmPFC ROI comprised the medial orbitofrontal cortex and gyrus rectus labels (Yoon et al., 2018; Neudert et al., 2023). All the anatomically defined masks were created by using WFU_pickatlas (Maldjian et al., 2003).

## Results

### Study population

We recruited 90 participants (n=45 cTBS group, n=45 Sham group) who completed the behavioural task. The behavioural sample consisted of 30 male and 60 female participants, of which 15 men and 30 women were assigned to each intervention group. The fMRI sample included 87 participants (44 cTBS, 43 Sham). There were no significant differences (p = 0.890) in the age distribution between the TMS group (mean = 24.9 years, SD = 4.43) and SHAM group (mean = 24.5 years, SD = 4.10). Median age was 21 (age range: 18-39) years. Of the subjects, 86 were right-handed (43 per group) and 4 were left-handed (2 per group). While blinding was sufficient in the Sham group (48% assumed their group assignment correctly, more subjects in the cTBS group (78%) believed correctly that they had been assigned to the cTBS group. The Sham group, as compared to cTBS group, was less vigilant during the task (group effect: χ^2^ = 7.19, p = 0.007). There were no differences between groups for clinical or neurocognitive parameters (see table 1).

**Table 1:**
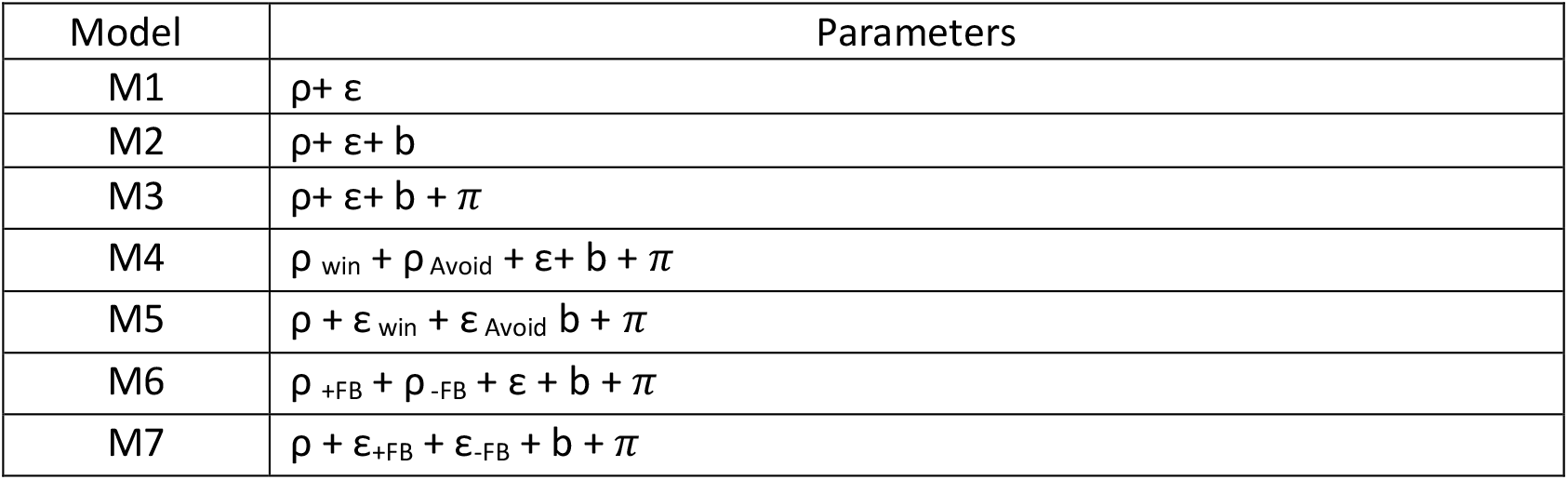
Models M1–M3, M5, and M7 use a single feedback-sensitivity parameter. M4 introduces context-specific feedback sensitivities, M5 context-specific learning rates, M6 valence-specific feedback sensitivities, and M7 valence-specific learning rates. All models from M2 onward include the Go-bias b, and models M3–M7 additionally include the motivational bias π. All models use separate model parameters for each block of the experiment.

**Table 1:** Group comparison of clinical parameters (STAI-G: State-Trait Anxiety Inventory Questionnaire, BDI: Beck Depression Inventory II, BIS: Barratt Impulsiveness Scale for impulsivity, OCI: Obsessive-Compulsive, ADHS-SB: ADHD scale for self-evaluation) and working memory tests. As the variables were not normally distributed, median and range are displayed and Mann-Whitney tests were used for group comparison.

|  | cTBS group<br>(n=45) median<br>(min-max) | Sham group (n=45)<br>median (min-max) | Group difference p<br>value | Effect size of group<br>difference (rang<br>biserial correlation) |
| --- | --- | --- | --- | --- |
| 80% active<br>motor<br>threshold (in<br>%) | 36 (28-55) | 50 (50-50) | < 0.001 | 0.91 |
| STAI-G | 33.0 (20-61) | 33.5 (21-58) | 0.7 | 0.37 |
| OCI | 4.0 (3-12) | 3.5 (3-14) | 0.1 | 0.1 |
| BIS | 12.0 (7-18) | 13.0 (9-18) | 0.6 | 0.03 |
| BDI | 2.0 (0-18) | 2.5 (0-17) | 0.9 | 0.01 |
| ADHS-SB | 7.0 (1-22) | 7.0 (1-22) | 0.97 | 0.0 |
| Digit-span<br>test level | 5 (4-8) | 5 (3-8) | 0.46 | 0.08 |
| Listening span<br>length | 3.5 (1-5.5) | 3.5 (2-6) | 0.41 | 0.09 |

### Behavioural results

#### Go-Tendency

The cTBS group showed overall fewer Go-responses compared to the Sham group (group effect: z = 2.8, *p* = 0.004, see figure 2a/b); this effect was independent of required action (group * action interaction: z = 0.5, *p* = 0.58). There were marginally significant interaction effects between group x valence (z = 1.7, p = 0.08) and group * action * valence (z = 1.7, *p* = 0.08). Both these trends of interaction effects resulted from a more pronounced reduction of go responses in win trials (post-hoc testing: z-ratio = 3.2, p = 0.001, Figure 2b). Group-independent task effects were similar as described previously (Swart et al., 2017; Scholz et al., 2022): A motivational bias was evident in both groups, indicated by more Go responses after Win relative to Avoid Punishment cues (valence effect: z = 8.7, *p* < 0.001). There was no main effect of block (*p* = 0.68) or treatment * block interaction effect (*p* = 0.58) on the Go-Tendency. There was a marginally significant interaction effect of valence * block (z = 1.9, *p* = 0.06). This resulted from fewer go responses during avoid trials in the first block compared to the second block. There were no other significant effects.

**Figure 2.**
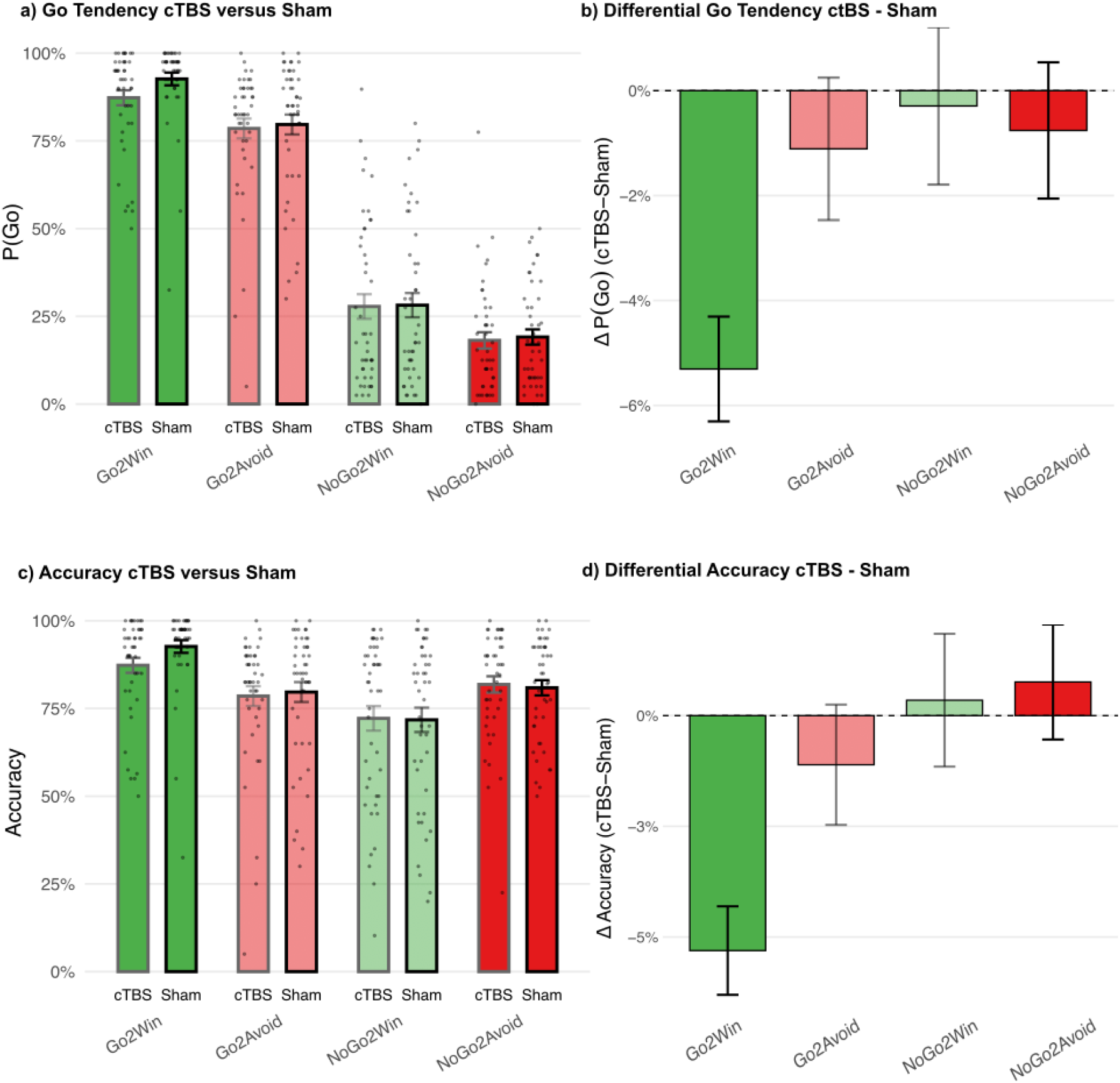
A) Proportion of Go-Responses (±SED) in the cTBS versus Sham group, across all trials, separated for each trial type. B) Difference of proportions of Go-Responses between cTBS and Sham group for each trial type. cTBS reduced the overall Go-proportion, with the biggest reduction in Go-to-Win trials. C) Mean (±SED) accuracy, i.e. proportion of correct responses, in the cTBS versus Sham group, across all trials. D) Difference of mean accuracy between cTBS and Sham group for each trial type. cTBS reduced accuracy in Go trials, especially in Go-to-Win trials.

#### Accuracy

The overall fewer Go responses in the cTBS group consequently led to a lower accuracy of the cTBS group in Go trials (group * action: z = 2.8, *p* = 0.004). There were marginally significant interaction effects between group x valence (z = 1.7, p = 0.08) and group * action * valence (z = 1.8, p = 0.07). Coherent with the results of the Go-models, these trends of interaction effects resulted from a reduced accuracy specifically in Go2Win trials (post-hoc testing: z-ratio = 2.5, p = 0.01, Figure 2d). The groups did not differ significantly in overall accuracy (group: z = 0.5, *p* = 0.6). In both groups, subjects had a significantly higher accuracy in the bias-congruent Go2Win trials, compared to the bias-incongruent NoGo2Win trials (action * valence interaction: z = 8.6, *p* < 0.001). Participants of both groups had a higher accuracy in Block 2 (block effect: z = 4.3, *p* < 0.001), indicating a training effect. This effect was driven by higher accuracies in block 2, especially in bias incongruent trial types (Go2Avoid and NoGo2Win, action * valence * block interaction: z = 1.9, p = 0.06). There was no other significant main effect or interaction.

#### Reaction Time

The cTBS group showed slower RTs relative to the Sham group (group: t = 2.0, p = 0.04), especially in NoGo trials (group * action effect: t = 2,3, p = 0.02). Overall, participants across groups also showed slower RT during avoid trials as compared to win trials, indicating a significant main effect of valence but no interaction with group (valence: t = 14.0, p < 0.001; group * valence effect: p = 0.12). Participants had faster reaction times in block 2 (block effect: t = 2.2, p = 0.03), this effect was driven by faster reaction times in block 2 in Go trials (action * block effect: z = 4.7, p < 0.001). There was no other significant main effect or interaction.

#### Computational modelling

Based on the model comparison, model M7, characterized by a single feedback-sensitivity parameter, separate learning rates for positive and negative feedback, and a single Go-Bias and Pavlovian-bias parameter, exhibited both the highest model frequency (0.63) and Protected exceedance probability (0.99) within the model space. Based on simulations, it also recovered the observed data well (Figure 3_f).

**Figure 3.**
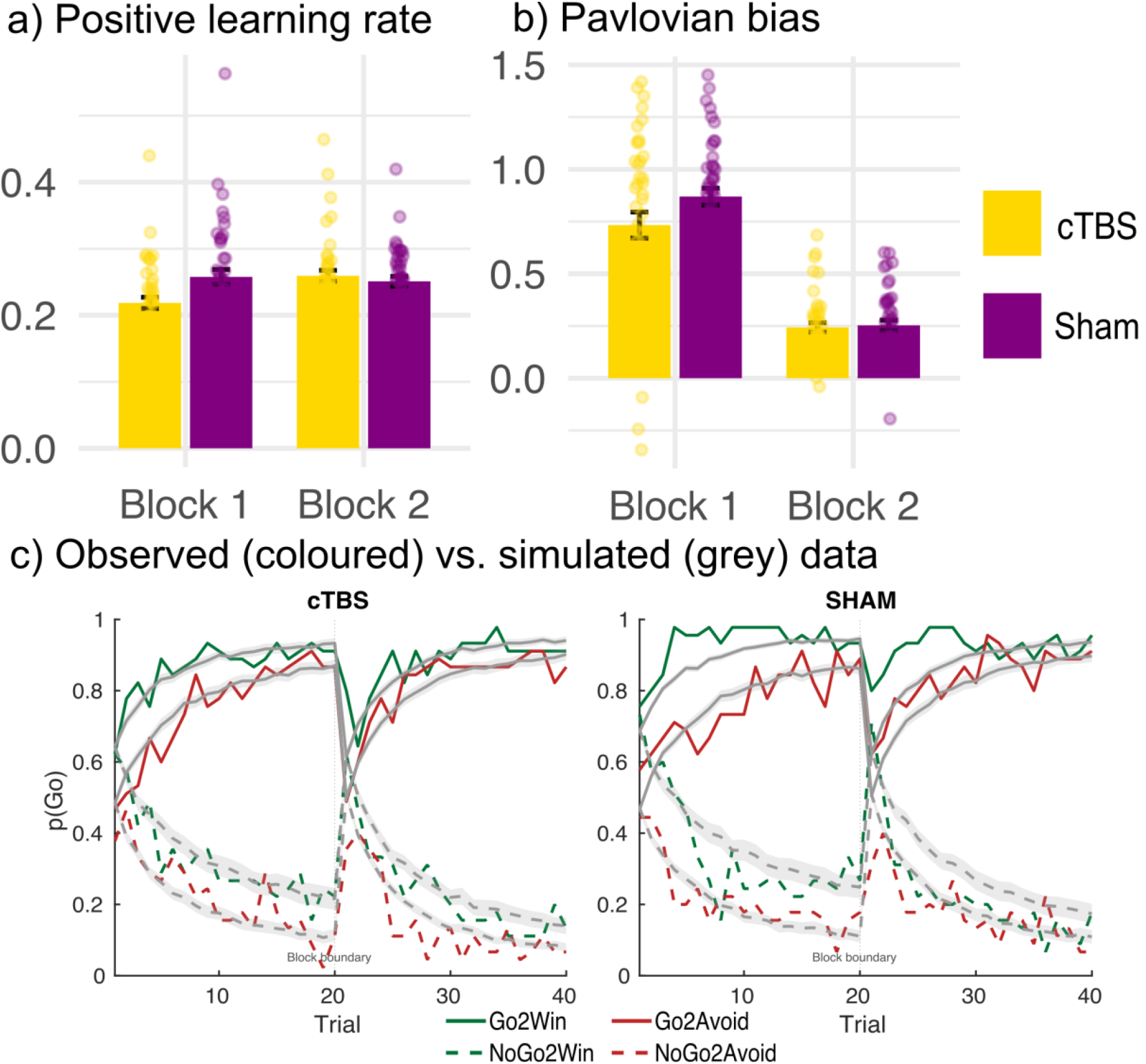
The positive learning rate parameter (a) was significantly reduced in the first block after cTBS. The Pavlovian Bias parameter (b), capturing motivational biases, was reduced after cTBS as a trend. The remaining parameters, reinforcement sensitivity, negative learning rate, and Go bias, were not significantly affected by cTBS, indicated by p values > 0.1 for all main effects of condition or interaction terms. Simulations (green and red lines representing cue valence) of our winning model 7 and the observed data (coloured lines) show that the decrease in positive learning rate after cTBS led to slower learning in the early Go2Win trials (c).

We subsequently examined parameter estimates of the winning model (M7) using linear mixed models (Parameter ∼ Group * Block + (1 | ID)): Reinforcement sensitivity increased in both groups during the second block relative to the first (t = 3.2, p = 0.002), while the Pavlovian Bias decreased in both groups during the second block (t = 14.1, p < 0.001). A significant group × block interaction emerged for the positive learning rate, driven by a lower learning rate in the cTBS as compared to the sham group, particularly during the first block (in block 1, t = 2.9, p = 0.005). Finally, there was a statistical trend toward an overall greater Pavlovian bias in the Sham versus TMS group across blocks (z = 1.8, p = 0.08). All remaining parameters showed no significant main effects or interactions (p > 0.1). Significant group comparisons are presented in Figure 3a/b.

#### MRI results

Whole-brain analyses revealed significant main effects across all participants for each regressor of interest. Model-derived choice probability correlated positively with activation primarily in the vmPFC (MNI: 0, 54, −10) and the left primary motor cortex (MNI: −32, −22, 50), see figure 4a. Button-press onset correlated positively with activation primarily in the frontal eye fields (MNI: 2, 30, 54), the right parietal cortex (MNI: 60, −38,46), and the right insula (MNI: 32, 20, −14). The reward-prediction error (RPE) signal was associated with significant activity in several regions, including the vmPFC (MNI: 8, 42, −6), the bilateral caudate nucleus (MNI: 8, 12, −8), and the occipital cortex (e.g. −36, −74, 34), see figure 4b. All reported clusters had a significance of p_FWE_ < 0.001.

**Figure 4.**
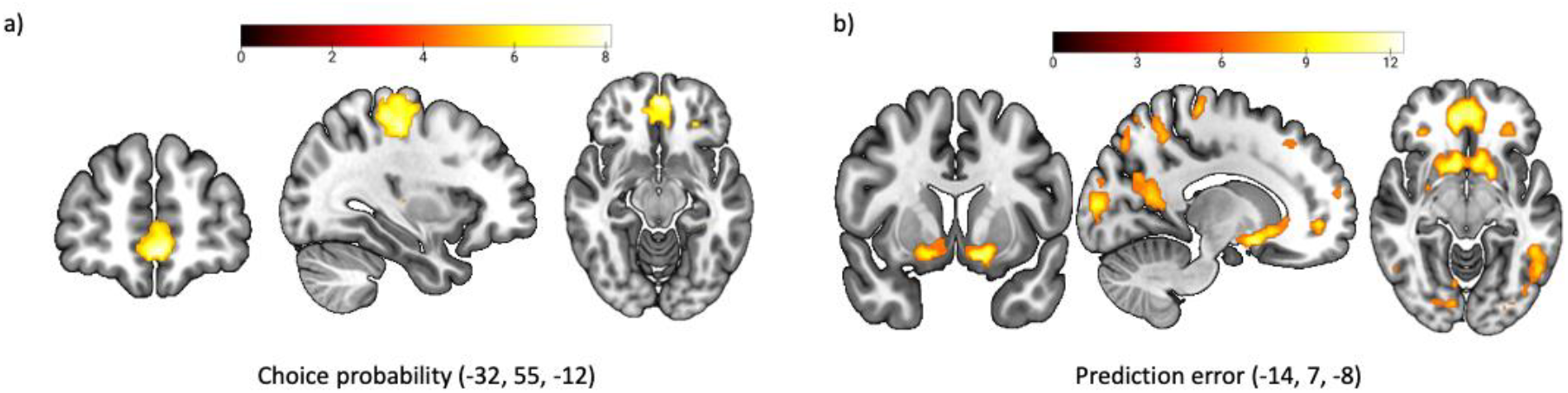
Main effects of the parametric modulator choice probability at cue onset (a) and prediction error at outcome onset (b). The effects are corrected for p_FWE_ < 0.001. Colourbar represents t-values.

A second-level contrast comparing the cTBS and sham group for the parametric RPE regressor revealed three foci with significantly different RPE-related activity (see figure 5): Within the anatomically defined mask of the vmPFC, heightened RPE responses were observed in the cTBS group (MNI = [10 32 –16]; 9 voxels; t = 4.19; pFWE = 0.030, peak-level SVC). SVC confined to the left striatum likewise indicated greater RPE-related activity in the putamen of the cTBS group (MNI = [–26 –8 –8]; 6 voxels; t = 4.12; pFWE = 0.030, peak-level SVC). The homologous effect in the right striatum did not reach significance (MNI = [20 –16 –20]; 10 voxels; t = 3.53; pFWE = 0.10). For completion, we report that at the whole brain level, a single cluster in the thalamus (MNI = [2 –20 10]; 238 voxels) showed stronger RPE signalling in the cTBS as compared to the sham group (t = 5.40; pFWE = 0.030, peak-level, whole-brain corrected, see figure 5c). No additional between-group differences survived correction for multiple comparisons in any other contrast or region of interest.

**Figure 5.**
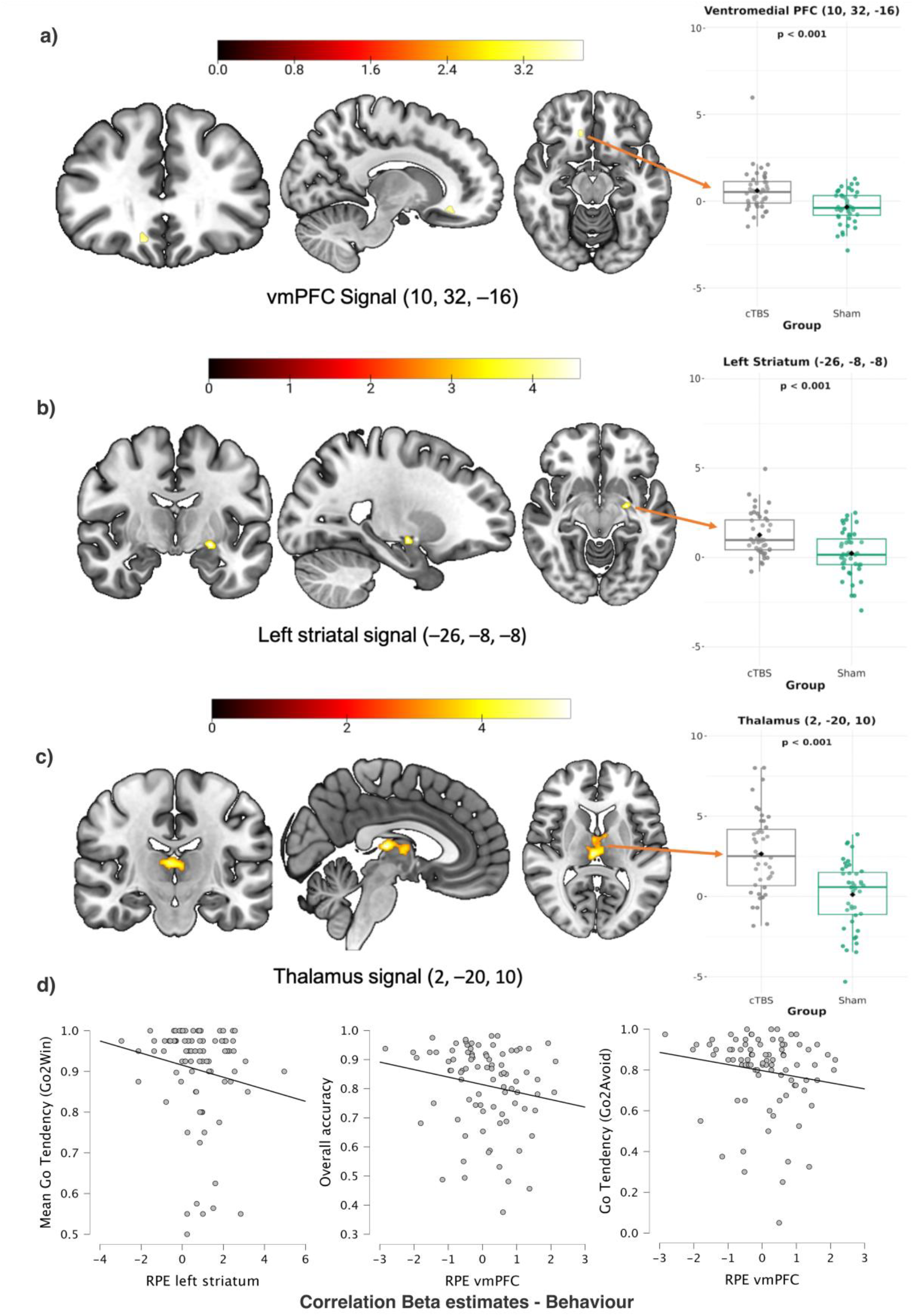
Statistical parametric maps illustrate significantly stronger BOLD responses to the Prediction-Error regressor after continuous theta-burst stimulation (cTBS) in (a) ventromedial prefrontal cortex (vmPFC), (b) left striatum, and (c) thalamus. The vmPFC and striatal peaks survive small-volume family-wise-error correction (p<0.05, SVC), whereas the thalamic cluster survives whole-brain FWE correction at p<0.05. Only the significant clusters are extracted for clearer visibility. The colour bar encodes t-values. Images are displayed in radiological convention (left = right). The bar plots on the right display the beta-estimates, extracted at each peak voxel of each of the significant clusters. Coloured dots represent individual subjects; the black diamond marks the group mean. Two-sample t-test p-values for the cTBS-versus-Sham comparison are printed above each panel. (d) Beta estimates of the striatum correlated with a lower Go-tendency in Go2Win trials, and vmPFC beta estimates correlated with an overall lower accuracy and a lower Go-tendency in Go2Avoid trials.

Additional fMRI analyses: To test whether cTBS over the vmPFC specifically modulated neural responses to cue valence (Win vs. Avoid / Go vs. NoGo) and outcome valence (positive vs. negative), we estimated two additional GLMs.

Model 1 included the following regressors: onsets of Go-to-Win, NoGo-to-Win, Go-to-Avoid, and NoGo-to-Avoid cues; button press; and outcome onset with model-derived prediction error (PE) as a parametric modulator. Results largely mirrored the main analysis, showing increased PE-related activity in the vmPFC (small-volume corrected, p = 0.02, T = 4.3, x = 10, y = 32, z = –16), in the thalamus (trend-level, p = 0.07, T = 5.12, x = 4, y = –20, z = 10), and in the left striatum (small-volume corrected, p = 0.02, T = 4.3, x = –26, y = –8, z = –8). No additional group-level differences emerged (p_min = 0.7).

To test for group differences in neural representations of positive and negative prediction error signals, model 2 included cue onsets; button press; positive outcomes (+100 after Win cues, 0 after Avoid cues), modelled with positive PE as a parametric modulator; and negative outcomes (0 after Win cues, –100 after Avoid cues), modelled with negative PE as a parametric modulator. This model revealed no significant group-level differences in neural activation (p_min = 0.6).

#### Correlations between prediction error signals and behaviour

To examine the relationship between neural prediction error signals and behavioural performance, we extracted individual beta estimates from the peak voxel of each of the three clusters showing significant group differences (thalamus, left striatum, and vmPFC). The prediction error signal in the left striatum correlated negatively with participants’ mean Go tendency and accuracy in Go2Win trials (r = –0.24, p = 0.02). The vmPFC prediction error signal correlated negatively with individuals’ mean Go tendency and accuracy in Go2Avoid trials (r = –0.23, p = 0.03), as well as with overall accuracy across all trial types (r = –0.23, p = 0.03, see Figure 5d). No other correlations reached significance (all p ≥ 0.07). We excluded one outlier from the correlation analysis.

## Discussion

In this study, transient disruption of vmPFC activity reduced overall Go responding, slowed reaction times, and selectively lowered the positive learning-rate parameter of the RL model, particularly during the first block. TMS also tended to weaken Pavlovian invigoration of Go responses for reward cues, indicated by a trend towards a reduced Pavlovian bias. On the neural level, cTBS amplified reward-prediction-error (RPE) signals in a vmPFC– mediodorsal thalamus-putamen network, while cue-value/choice-probability signals at cue onset remained unchanged. Stronger RPE-signals in the TMS group were correlated with lower go response and reduced performance.

We interpret the stronger RPE signals as a signature of higher uncertainty (lower precision). When value beliefs are imprecise, outcomes become less predictable, reflected in larger RPEs. This uncertainty account can link our behavioural and neural findings. The reduction of the positive learning rate after cTBS implies slower acquisition of action–reward associations, which may counteract the TMS effect on RPEs, in particular in a task with stable reward-outcome contingencies. Consistent with this, greater PE-BOLD correlated with poorer performance (left striatum PE with lower Go tendency in Go2Win trials; vmPFC PE with lower overall accuracy). Consistent with this, enhancement of neural prediction-error signalling following vmPFC inhibition has previously been demonstrated (Starkweather et al., 2018).

The altered activation pattern of RPEs is part of a well-established cortico-subcortical network involved in reward learning and decision-making. The vmPFC has dense anatomical projections to subcortical targets such as the nucleus accumbens and dorsal striatum (Vertes, 2004; Gabbott et al., 2005; Kim et al., 2017), forming a cortico-striatal loop that is in close, reciprocal communication with the MD thalamus (Gabbott et al., 2005; Haber and Knutson, 2010; Mitchell, 2015; Peters et al., 2016). Electrophysiological studies have shown that prefrontal cortical neurons transiently encode cue values and subsequently activate MD thalamic neurons, which then amplify and sustain cue-value representations until a motor decision is made (Preuss and Goldman-Rakic, 1987; Schmitt et al., 2017). Lesions anywhere in this circuit impair reward learning, prolonging decision latencies and leading to inflexible behaviour and a weaker exploitation of recent feedback (St Onge et al., 2012; Narayanan et al., 2013; Browning et al., 2015). Dysfunction in this circuitry has also been associated with impaired cognitive performance, a characteristic feature of psychiatric conditions such as schizophrenia (Parnaudeau et al., 2013). Crucially, inhibiting the medial prefrontal cortex in rodents paradoxically increases MD firing in the thalamus (Schmitt et al., 2017; Li et al., 2024) by refining the tuning of individual cortical neurons and recruiting previously inactive neurons. These dynamics may represent an attention-regulating mechanism that sustains cortical activity when initial prefrontal signals diminish over time and could explain why our results do not show any weaker neural activation in the vmPFC after cTBS. Strengthening the cortico-thalamic drive, though eventually less efficient, preserves cue-value coding under uncertainty by boosting instrumental, action-outcome-associated learning (Corbit et al., 2003; Schmitt et al., 2017; Rikhye et al., 2018; Wang et al., 2023). This shift could explain the observed trend towards a lower Pavlovian bias modelling parameter, as hypothesised in previous studies (Algermissen et al., 2024).

This points to a specific role of the vmPFC in the context of motivational learning and decision-making: The vmPFC mediates trial-by-trial behavioural adaptation with positive-valenced signals to midbrain dopamine neurons via ventral striatal relays (Haber and Calzavara, 2009; Haber and Behrens, 2014; Ferenczi et al., 2016; Gentry and Roesch, 2018). Reward-predictive cues subsequently elicit phasic dopamine bursts that bias the D1-expressing “direct” striatal pathway toward action (Kravitz et al., 2012; Collins and Frank, 2014; Coddington and Dudman, 2018). PET studies show that higher baseline striatal dopamine predicts stronger positive learning rates and greater reward-driven Go tendencies in humans (de Boer et al., 2019). They also indicate that individuals with higher working-memory and striatal baseline dopamine levels exhibit a stronger Pavlovian bias (Chen et al., 2023b). A global dopamine increase due to levodopa administration reduced Pavlovian bias, possibly by improving prefrontal control (Guitart-Masip et al., 2014b), invigorated motor responding (Guitart-Masip et al., 2012a), and increased learning rates (Chowdhury et al., 2013). In addition, enhancing catecholamine levels using methylphenidate increased the Pavlovian bias in individuals with higher working memory but decreased the bias in individuals with low working memory (Swart et al., 2017). Therefore, vmPFC cTBS might have weakened the corticodopaminergic ‘go’ drive, reducing the positive learning rate. This, in turn, would have dampened D1-mediated action invigoration, manifesting behaviourally as fewer and slower go responses.

One plausible driver of the observed slower Go responses after cTBS could be an evidence-accumulation bottleneck, characterized by a higher decision threshold, emerging when the vmPFC can no longer rapidly supply context-appropriate value signals. Drift-diffusion analyses have demonstrated that vmPFC lesions elevate choice noise and prolong evidence integration (Yu et al., 2022). Similarly, trial-to-trial instability in vmPFC value coding predicts delayed and inconsistent decision-making (Kurtz-David et al., 2019; Ciaramelli et al., 2021; Moneta et al., 2023). These behavioural patterns have been computationally described using RL models with a “softmax temperature” parameter, reflecting the precision or noisiness of value-based decision-making. Higher softmax temperatures indicate noisier, less precise choices. Notably, magnetic-resonance spectroscopy has linked higher cortical GABA levels, whose inhibitory effects resemble those of cTBS, to elevated softmax temperatures, slower value accumulation, and thus slower decision-making. Conversely, increased cortical glutamate levels correlated with lower softmax temperatures and faster decision dynamics (Jocham et al., 2012; Kaiser et al., 2021). Collectively, these findings suggest that the vmPFC plays a critical role in rapid and consistent decision-making by filtering out irrelevant sensory information and stabilizing value representations (Moneta et al., 2023; Broschard et al., 2024).

Although vmPFC disruption resulted in less go behavior in our task, the fixed contingencies enabled the preservation of a stable policy representation by distributed networks, as demonstrated by the unaltered cue-onset choice-probability signal. This finding supports the hypothesis that value-based decision-making involves a distributed network that extends beyond the vmPFC to encompass other prefrontal areas (Rushworth et al., 2011; Guitart-Masip et al., 2012b; Badre, 2021), as well as the cerebellum and striatum (Zyuzin et al., 2023). Within this distributed circuitry, the vmPFC primarily integrates recent outcome signals to rapidly update stimulus values, rather than maintaining stable, long-term action-value representations. For instance, the dlPFC has been linked to the maintenance of goal-relevant information and the support of decision-making processes in the event of impaired vmPFC function (Hunt and Hayden, 2017; Tomov et al., 2020). In support of this interpretation, previous studies have shown minimal impairment in deterministic cue–reward learning tasks following vmPFC lesions (Fellows and Farah, 2003, 2005). Consequently, disruption to the vmPFC may selectively impair the rapid, feedback-driven learning of representations of choice probability without fundamentally compromising them. While our primary focus was on implementing and evaluating the causal impact of the vmPFC disruption, additional modelling analyses, like separating signed and unsigned prediction errors (Algermissen et al., 2024), could help further elucidate the role of the vmPFC in learning and behavioural control.

## Limitations

First, the observed lateralisation of small-volume-corrected BOLD effects in the vmPFC and putamen, with bilateral neural activation confined to the thalamic RPE increases, may either reflect genuine hemispheric specialisation or limited statistical sensitivity. Future studies using larger sample sizes could clarify this potential lateralisation, shedding additional light on hemisphere-specific contributions to Pavlovian-instrumental interactions. Second, while we operated under the established assumption that continuous theta-burst stimulation (cTBS) reliably suppresses cortical excitability for approximately 30–60 minutes, in a non-motor area such as the vmPFC, we could not record markers such as motor-evoked potentials to directly confirm this. Incorporating indirect markers of cortical excitability, such as MR spectroscopy, in future research could precisely define stimulation depth and temporal dynamics, further refining the interpretation and reproducibility of cTBS-related findings. Third, stimulation delivered at Fpz with a figure-of-eight coil inherently involves adjacent medial and orbitofrontal cortical areas. While this anatomical proximity limits the ability to target vmPFC microcircuits with high precision using TMS, cTBS remains one of the most effective non-invasive tools for spatially restricted modulation of prefrontal cortex function in humans (Lowe et al., 2018). Fourth, our results show that participants in the cTBS group assumed their group assignment more accurately. In future studies, including an active sham group, stimulating another cortical region could rule out possible deblinding effects.

## Conclusions

Across species, the vmPFC has been shown to map and flexibly update reward expectations (Tremblay and Schultz, 1999; Lopatina et al., 2016) while also guiding adaptive choice (Izquierdo et al., 2004). However, valuation processes are also robustly distributed across multiple cortical and subcortical structures (Hunt and Hayden, 2017; Hiser and Koenigs, 2018). The current findings refine this conceptual framework by showing that transient vmPFC inhibition triggers a shift from appetitively biased Pavlovian mechanisms toward a thalamus-vmPFC and dorsal-striatum-centered instrumental learning strategy, resulting in longer RTs and with an early cost in Go invigoration and positive-feedback learning. Amplified RPEs in the thalamus, putamen, and vmPFC may reflect elevated uncertainty and hence stronger outcome surprise signals. The unchanged cue-onset choice probabilities in the cTBS group indicate that dorsal fronto-striatal circuits effectively maintain stable policy representations, even when vmPFC-dependent value updating is disrupted. Taken together, these data suggest that the vmPFC contributes critically to the stabilisation of preferences, minimisation of behavioural variability (Moneta et al., 2023), and the rapid, adaptive updating of reward-guided behaviour.

## References

Algermissen J, Swart JC, Scheeringa R, Cools R, den Ouden HEM (2021) Striatal BOLD and midfrontal theta power express motivation for action. Cerebral Cortex 32:2924–2942.

Algermissen J, Swart JC, Scheeringa R, Cools R, den Ouden HEM (2024) Prefrontal signals precede striatal signals for biased credit assignment in motivational learning biases. Nature Communications 15:19.

Ashburner J, Friston KJ (2005) Unified segmentation. neuroimage 26:839–851.

Badre D (2021) Navigating the recurrent perplexity of prefrontal function. Brain 145:406–407.

Barr DJ, Levy R, Scheepers C, Tily HJ (2013) Random effects structure for confirmatory hypothesis testing: Keep it maximal. J Mem Lang 68.

Bartra O, McGuire JT, Kable JW (2013) The valuation system: a coordinate-based meta-analysis of BOLD fMRI experiments examining neural correlates of subjective value. Neuroimage 76:412–427.

Blackburn HL, Benton AL (1957) Revised administration and scoring of the digit span test. Journal of consulting psychology 21:139.

Broschard MB, Turner BM, Tranel D, Freeman JH (2024) Dissociable Roles of the Dorsolateral and Ventromedial Prefrontal Cortex in Human Categorization. The Journal of Neuroscience 44:e2343232024.

Browning PG, Chakraborty S, Mitchell AS (2015) Evidence for mediodorsal thalamus and prefrontal cortex interactions during cognition in macaques. Cerebral cortex 25:4519–4534.

Cavanagh JF, Eisenberg I, Guitart-Masip M, Huys Q, Frank MJ (2013) Frontal theta overrides pavlovian learning biases. J Neurosci 33:8541–8548.

Chen K, Garbusow M, Sebold M, Kuitunen-Paul S, Smolka MN, Huys QJM, Zimmermann US, Schlagenhauf F, Heinz A (2023a) Alcohol Approach Bias Is Associated With Both Behavioral and Neural Pavlovian-to-Instrumental Transfer Effects in Alcohol-Dependent Patients. Biol Psychiatry Glob Open Sci 3:443–450.

Chen P, Geurts DEM, Määttä JI, van den Bosch R, Hofmans L, Papadopetraki D, den Ouden H, Cools R (2023b) Effect of striatal dopamine on Pavlovian bias. A large [^18^F]-DOPA PET study. Behavioral Neuroscience 137:184–195.

Chowdhury R, Guitart-Masip M, Lambert C, Dayan P, Huys Q, Düzel E, Dolan RJ (2013) Dopamine restores reward prediction errors in old age. Nature Neuroscience 16:648–653.

Ciaramelli E, De Luca F, Kwan D, Mok J, Bianconi F, Knyagnytska V, Craver C, Green L, Myerson J, Rosenbaum RS (2021) The role of ventromedial prefrontal cortex in reward valuation and future thinking during intertemporal choice. eLife 10:e67387.

Coddington LT, Dudman JT (2018) The timing of action determines reward prediction signals in identified midbrain dopamine neurons. Nature Neuroscience 21:1563–1573.

Collins AG, Frank MJ (2014) Opponent actor learning (OpAL): modeling interactive effects of striatal dopamine on reinforcement learning and choice incentive. Psychological review 121:337.

Corbit LH, Muir JL, Balleine BW (2003) Lesions of mediodorsal thalamus and anterior thalamic nuclei produce dissociable effects on instrumental conditioning in rats. The European journal of neuroscience 18:1286–1294.

Daneman M, Carpenter PA (1980) Individual differences in working memory and reading. Journal of Verbal Learning and Verbal Behavior 19:450–466.

Dantas AM, Sack AT, Bruggen E, Jiao P, Schuhmann T (2023) The functional relevance of right DLPFC and VMPFC in risk-taking behavior. Cortex 159:64–74.

Daw ND, Gershman SJ, Seymour B, Dayan P, Dolan RJ (2011) Model-based influences on humans’ choices and striatal prediction errors. Neuron 69:1204–1215.

de Boer L, Axelsson J, Chowdhury R, Riklund K, Dolan RJ, Nyberg L, Bäckman L, Guitart-Masip M (2019) Dorsal striatal dopamine D1 receptor availability predicts an instrumental bias in action learning. Proceedings of the National Academy of Sciences 116:261–270.

Fellows LK, Farah MJ (2003) Ventromedial frontal cortex mediates affective shifting in humans: evidence from a reversal learning paradigm. Brain 126:1830–1837.

Fellows LK, Farah MJ (2005) Different underlying impairments in decision-making following ventromedial and dorsolateral frontal lobe damage in humans. Cereb Cortex 15:58–63.

Ferenczi EA, Zalocusky KA, Liston C, Grosenick L, Warden MR, Amatya D, Katovich K, Mehta H, Patenaude B, Ramakrishnan C (2016) Prefrontal cortical regulation of brainwide circuit dynamics and reward-related behavior. Science 351:aac9698.

Gabbott PL, Warner TA, Jays PR, Salway P, Busby SJ (2005) Prefrontal cortex in the rat: projections to subcortical autonomic, motor, and limbic centers. J Comp Neurol 492:145–177.

Gentry RN, Roesch MR (2018) Neural Activity in Ventral Medial Prefrontal Cortex Is Modulated More Before Approach Than Avoidance During Reinforced and Extinction Trial Blocks. J Neurosci 38:4584–4597.

Gershman SJ, Guitart-Masip M, Cavanagh JF (2021) Neural signatures of arbitration between Pavlovian and instrumental action selection. PLoS computational biology 17:e1008553.

Gönner S, Leonhart R, Ecker W (2007) Das Zwangsinventar OCI-R-die deutsche version des obsessivecompulsive inventory-revised. PPmP-Psychotherapie· Psychosomatik· Medizinische Psychologie 57:395–404.

Guitart-Masip M, Duzel E, Dolan R, Dayan P (2014a) Action versus valence in decision making. Trends Cogn Sci 18:194–202.

Guitart-Masip M, Chowdhury R, Sharot T, Dayan P, Duzel E, Dolan RJ (2012a) Action controls dopaminergic enhancement of reward representations. Proceedings of the National Academy of Sciences 109:7511–7516.

Guitart-Masip M, Huys QJ, Fuentemilla L, Dayan P, Duzel E, Dolan RJ (2012b) Go and no-go learning in reward and punishment: interactions between affect and effect. Neuroimage 62:154–166.

Guitart-Masip M, Fuentemilla L, Bach DR, Huys QJ, Dayan P, Dolan RJ, Duzel E (2011) Action dominates valence in anticipatory representations in the human striatum and dopaminergic midbrain. J Neurosci 31:7867–7875.

Guitart-Masip M, Economides M, Huys QJ, Frank MJ, Chowdhury R, Duzel E, Dayan P, Dolan RJ (2014b) Differential, but not opponent, effects of L-DOPA and citalopram on action learning with reward and punishment. Psychopharmacology 231:955–966.

Haber SN, Calzavara R (2009) The cortico-basal ganglia integrative network: the role of the thalamus. Brain research bulletin 78:69–74.

Haber SN, Knutson B (2010) The reward circuit: linking primate anatomy and human imaging. Neuropsychopharmacology 35:4–26.

Haber SN, Behrens TE (2014) The neural network underlying incentive-based learning: implications for interpreting circuit disruptions in psychiatric disorders. Neuron 83:1019–1039.

Herrmann MJ, Katzorke A, Busch Y, Gromer D, Polak T, Pauli P, Deckert J (2017) Medial prefrontal cortex stimulation accelerates therapy response of exposure therapy in acrophobia. Brain stimulation 10:291–297.

Hiser J, Koenigs M (2018) The Multifaceted Role of the Ventromedial Prefrontal Cortex in Emotion, Decision Making, Social Cognition, and Psychopathology. Biol Psychiatry 83:638–647.

Huang YZ, Edwards MJ, Rounis E, Bhatia KP, Rothwell JC (2005) Theta burst stimulation of the human motor cortex. Neuron 45:201–206.

Hunt LT, Hayden BY (2017) A distributed, hierarchical and recurrent framework for reward-based choice. Nat Rev Neurosci 18:172–182.

Huys QJ, Lally N, Faulkner P, Eshel N, Seifritz E, Gershman SJ, Dayan P, Roiser JP (2015) Interplay of approximate planning strategies. Proc Natl Acad Sci U S A 112:3098–3103.

Izquierdo A, Suda RK, Murray EA (2004) Bilateral orbital prefrontal cortex lesions in rhesus monkeys disrupt choices guided by both reward value and reward contingency. J Neurosci 24:7540–7548.

Jackson-Koku G (2016) Beck depression inventory. Occupational medicine 66:174–175.

Jocham G, Hunt LT, Near J, Behrens TE (2012) A mechanism for value-guided choice based on the excitation-inhibition balance in prefrontal cortex. Nat Neurosci 15:960–961.

Kaiser LF, Gruendler TOJ, Speck O, Luettgau L, Jocham G (2021) Dissociable roles of cortical excitation-inhibition balance during patch-leaving versus value-guided decisions. Nature Communications 12:904.

Kass RE, Raftery AE (1995) Bayes factors. Journal of the american statistical association 90:773–795.

Kim CK, Ye L, Jennings JH, Pichamoorthy N, Tang DD, Yoo A-CW, Ramakrishnan C, Deisseroth K (2017) Molecular and circuit-dynamical identification of top-down neural mechanisms for restraint of reward seeking. Cell 170:1013-1027. e1014.

Kravitz AV, Tye LD, Kreitzer AC (2012) Distinct roles for direct and indirect pathway striatal neurons in reinforcement. Nature neuroscience 15:816–818.

Kurtz-David V, Persitz D, Webb R, Levy DJ (2019) The neural computation of inconsistent choice behavior. Nat Commun 10:1583.

Li F, Zheng X, Wang H, Meng L, Chen M, Hui Y, Liu D, Li Y, Xie K, Zhang J, Guo G (2024) Mediodorsal thalamus projection to medial prefrontal cortical mediates social defeat stress-induced depression-like behaviors. Neuropsychopharmacology 49:1318–1329.

Lo S, Andrews S (2015) To transform or not to transform: using generalized linear mixed models to analyse reaction time data. Frontiers in Psychology 6.

Lopatina N, McDannald MA, Styer CV, Peterson JF, Sadacca BF, Cheer JF, Schoenbaum G (2016) Medial Orbitofrontal Neurons Preferentially Signal Cues Predicting Changes in Reward during Unblocking. J Neurosci 36:8416–8424.

Lowe CJ, Manocchio F, Safati AB, Hall PA (2018) The effects of theta burst stimulation (TBS) targeting the prefrontal cortex on executive functioning: A systematic review and meta-analysis. Neuropsychologia 111:344–359.

Maia TV, Frank MJ (2011) From reinforcement learning models to psychiatric and neurological disorders. Nat Neurosci 14:154–162.

Maldjian JA, Laurienti PJ, Kraft RA, Burdette JH (2003) An automated method for neuroanatomic and cytoarchitectonic atlas-based interrogation of fMRI data sets. Neuroimage 19:1233–1239.

Manuel AL, Murray NWG, Piguet O (2019) Transcranial direct current stimulation (tDCS) over vmPFC modulates interactions between reward and emotion in delay discounting. Scientific Reports 9:18735.

Messimeris D, Levy R, Le Bouc R (2023) Economic and social values in the brain: evidence from lesions to the human ventromedial prefrontal cortex. Front Neurol 14:1198262.

Mitchell AS (2015) The mediodorsal thalamus as a higher order thalamic relay nucleus important for learning and decision-making. Neuroscience and biobehavioral reviews 54:76–88.

Moneta N, Garvert MM, Heekeren HR, Schuck NW (2023) Task state representations in vmPFC mediate relevant and irrelevant value signals and their behavioral influence. Nature Communications 14:3156.

Narayanan NS, Cavanagh JF, Frank MJ, Laubach M (2013) Common medial frontal mechanisms of adaptive control in humans and rodents. Nat Neurosci 16:1888–1895.

Neudert MK, Schäfer A, Zehtner RI, Fricke S, Seinsche RJ, Kruse O, Stark R, Hermann A (2023) Behavioral pattern separation is associated with neural and electrodermal correlates of context-dependent fear conditioning. Sci Rep 13:5577.

Odorfer TM, Homola GA, Reich MM, Volkmann J, Zeller D (2019) Increased Finger-Tapping Related Cerebellar Activation in Cervical Dystonia, Enhanced by Transcranial Stimulation: An Indicator of Compensation? Frontiers in Neurology 10.

Parnaudeau S, O’neill P-K, Bolkan SS, Ward RD, Abbas AI, Roth BL, Balsam PD, Gordon JA, Kellendonk C (2013) Inhibition of mediodorsal thalamus disrupts thalamofrontal connectivity and cognition. Neuron 77:1151–1162.

Peng Z, He L, Wen R, Verguts T, Seger CA, Chen Q (2022) Obsessive-compulsive disorder is characterized by decreased Pavlovian influence on instrumental behavior. PLoS Comput Biol 18:e1009945.

Penny WD, Friston KJ, Ashburner JT, Kiebel SJ, Nichols TE (2011) Statistical parametric mapping: the analysis of functional brain images: Elsevier.

Pershin I, Candrian G, Münger M, Baschera GM, Rostami M, Eich D, Müller A (2023) Vigilance described by the time-on-task effect in EEG activity during a cued Go/NoGo task. Int J Psychophysiol 183:92–102.

Peters SK, Dunlop K, Downar J (2016) Cortico-Striatal-Thalamic Loop Circuits of the Salience Network: A Central Pathway in Psychiatric Disease and Treatment. Front Syst Neurosci 10:104.

Piray P, Dezfouli A, Heskes T, Frank MJ, Daw ND (2019) Hierarchical Bayesian inference for concurrent model fitting and comparison for group studies. PLoS computational biology 15:e1007043.

Preuss TM, Goldman-Rakic PS (1987) Crossed corticothalamic and thalamocortical connections of macaque prefrontal cortex. Journal of Comparative Neurology 257:269–281.

Raab HA, Hartley CA (2020) Adolescents exhibit reduced Pavlovian biases on instrumental learning. Scientific Reports 10:15770.

Rangel A, Camerer C, Montague PR (2008) A framework for studying the neurobiology of value-based decision making. Nature Reviews Neuroscience 9:545–556.

Rescorla R, Wagner A (1972) A theory of Pavlovian conditioning: Variations in the effectiveness of reinforcement and nonreinforcement. Classical conditioning II: Current research and theory:64–99.

Rescorla RA, Solomon RL (1967) Two-process learning theory: Relationships between Pavlovian conditioning and instrumental learning. Psychol Rev 74:151–182.

Retz W, Retz-Junginger P, Römer K, Rösler M (2013) Standardisierte Skalen zur strukturierten Diagnostik der ADHS im Erwachsenenalter. Fortschritte der Neurologie· Psychiatrie 81:381–389.

Rikhye RV, Gilra A, Halassa MM (2018) Thalamic regulation of switching between cortical representations enables cognitive flexibility. Nat Neurosci 21:1753–1763.

Rushworth MF, Noonan MP, Boorman ED, Walton ME, Behrens TE (2011) Frontal cortex and reward-guided learning and decision-making. Neuron 70:1054–1069.

Schmitt LI, Wimmer RD, Nakajima M, Happ M, Mofakham S, Halassa MM (2017) Thalamic amplification of cortical connectivity sustains attentional control. Nature 545:219–223.

Scholz V, Waltmann M, Herzog N, Horstmann A, Deserno L (2024) Decrease in decision noise from adolescence into adulthood mediates an increase in more sophisticated choice behaviors and performance gain. PLoS Biology 22:e3002877.

Scholz V, Algermissen J, Kandroodi MR, Gillan C, den Ouden H (2025) Adaptive suppression of motivational biases increases with mood/anxiety traits.

Scholz V, Hook RW, Kandroodi MR, Algermissen J, Ioannidis K, Christmas D, Valle S, Robbins TW, Grant JE, Chamberlain SR, den Ouden HEM (2022) Cortical dopamine reduces the impact of motivational biases governing automated behaviour. Neuropsychopharmacology 47:1503–1512.

St Onge JR, Stopper CM, Zahm DS, Floresco SB (2012) Separate prefrontal-subcortical circuits mediate different components of risk-based decision making. J Neurosci 32:2886–2899.

Stanford MS, Mathias CW, Dougherty DM, Lake SL, Anderson NE, Patton JH (2009) Fifty years of the Barratt Impulsiveness Scale: An update and review. Personality and individual differences 47:385–395.

Starkweather CK, Gershman SJ, Uchida N (2018) The Medial Prefrontal Cortex Shapes Dopamine Reward Prediction Errors under State Uncertainty. Neuron 98:616-629.e616.

Stephan KE, Penny WD, Daunizeau J, Moran RJ, Friston KJ (2009) Bayesian model selection for group studies. Neuroimage 46:1004–1017.

Swart JC, Frank MJ, Maatta JI, Jensen O, Cools R, den Ouden HEM (2018) Frontal network dynamics reflect neurocomputational mechanisms for reducing maladaptive biases in motivated action. PLoS Biol 16:e2005979.

Swart JC, Frobose MI, Cook JL, Geurts DE, Frank MJ, Cools R, den Ouden HE (2017) Catecholaminergic challenge uncovers distinct Pavlovian and instrumental mechanisms of motivated (in)action. Elife 6.

Tenenbaum G, Furst D, Weingarten G (1985) A statistical reevaluation of the STAI anxiety questionnaire. Journal of clinical psychology 41:239–244.

Tombaugh TN (2004) Trail Making Test A and B: normative data stratified by age and education. Archives of clinical neuropsychology 19:203–214.

Tomov MS, Truong VQ, Hundia RA, Gershman SJ (2020) Dissociable neural correlates of uncertainty underlie different exploration strategies. Nature Communications 11:2371.

Tremblay L, Schultz W (1999) Relative reward preference in primate orbitofrontal cortex. Nature 398:704–708.

Tzourio-Mazoyer N, Landeau B, Papathanassiou D, Crivello F, Etard O, Delcroix N, Mazoyer B, Joliot M (2002) Automated Anatomical Labeling of Activations in SPM Using a Macroscopic Anatomical Parcellation of the MNI MRI Single-Subject Brain. NeuroImage 15:273–289.

van Nuland AJ, Helmich RC, Dirkx MF, Zach H, Toni I, Cools R, den Ouden HEM (2020) Effects of dopamine on reinforcement learning in Parkinson’s disease depend on motor phenotype. Brain 143:3422–3434.

van Wouwe NC, van den Wildenberg WPM, Ridderinkhof KR, Claassen DO, Neimat JS, Wylie SA (2015) Easy to learn, hard to suppress: The impact of learned stimulus–outcome associations on subsequent action control. Brain and Cognition 101:17–34.

Vertes RP (2004) Differential projections of the infralimbic and prelimbic cortex in the rat. Synapse 51:32–58.

Wang BA, Drammis S, Hummos A, Halassa MM, Pleger B (2023) Modulation of prefrontal couplings by prior belief-related responses in ventromedial prefrontal cortex. Front Neurosci 17:1278096.

Wischnewski M, Schutter DJ (2015) Efficacy and time course of theta burst stimulation in healthy humans. Brain stimulation 8:685–692.

Wittchen H-U, Zaudig M, Fydrich T (1997) Skid. Strukturiertes klinisches interview für DSM-IV. Achse I und II. Handanweisung.

Wittmann MK, Trudel N, Trier HA, Klein-Flügge MC, Sel A, Verhagen L, Rushworth MFS (2021) Causal manipulation of self-other mergence in the dorsomedial prefrontal cortex. Neuron 109:2353-2361.e2311.

Yoon L, Somerville LH, Kim H (2018) Development of MPFC function mediates shifts in self-protective behavior provoked by social feedback. Nature Communications 9:3086.

Yu LQ, Dana J, Kable JW (2022) Individuals with ventromedial frontal damage display unstable but transitive preferences during decision making. Nat Commun 13:4758.

Zhao D, Li Y, Liu T, Voon V, Yuan T-F (2020) Twice-Daily Theta Burst Stimulation of the Dorsolateral Prefrontal Cortex Reduces Methamphetamine Craving: A Pilot Study. Frontiers in Neuroscience Volume 14 −2020.

Zyuzin J, Combs D, Melrose J, Kodaverdian N, Leather C, Carrillo JD, Monterosso JR, Brocas I (2023) The neural correlates of value representation: From single items to bundles. Human brain mapping 44:1476–1495.

